# Bioluminescence dynamics in single germinating bacterial spores reveal metabolic heterogeneity

**DOI:** 10.1101/2020.01.14.906875

**Authors:** Zak Frentz, Jonathan Dworkin

## Abstract

Spore-forming bacteria modulate their metabolic rate by over 5 orders of magnitude as they transition between dormant spores and vegetative cells, and thus represent an extreme case of phenotypic variation. During environmental changes in nutrient availability, clonal populations of spore-forming bacteria exhibit individual differences in cell fate, timing of phenotypic transitions, and gene expression. One potential source of this variability is metabolic heterogeneity, but this has not yet been measured, as existing single-cell methods are not easily applicable to spores due to their small size and strong autofluorescence. Here, we use the bacterial bioluminescence system and a highly sensitive microscope to measure metabolic dynamics in thousands of *B. subtilis* spores as they germinate. We observe and quantitate large variations in the bioluminescence dynamics across individual spores that can be decomposed into contributions from variability in germination timing, the amount of endogenously produced luminescence substrate, and the intracellular reducing power. This work shows that quantitative measurement of spore metabolism is possible and thus it opens venues for future study of the thermodynamic nature of dormant states.

## Introduction

Some bacteria respond to nutrient depletion by differentiating from growing, vegetative cells into spores that can survive extreme temperatures, radiation, and long periods without nutrients, while retaining the ability to germinate and reinitiate growth in the presence of sufficient nutrients [1-3]. One of the most remarkable aspects of spore-forming bacteria is the extreme reduction of their metabolic rate, by at least 5 orders of magnitude, during sporulation [4]. Metabolic dormancy and resistance allow spores to survive for long times in harsh environments, which typically fluctuate. Dynamically sensing the environment requires energy expenditure [5], and thus is likely difficult for spores given their low or nonexistent metabolism. Therefore, sporulating populations must adapt to fluctuating environments through other strategies, such as phenotypic diversification, or “bet-hedging” [6, 7].

Significant heterogeneity is observed in populations during both the sporulation and germination transitions. For example, after nutrient limitation, only a portion of vegetative cells commit to sporulation, while other cells either lyse or survive in the vegetative state by using secondary metabolites [8]. The time at which a vegetative cell sporulates varies significantly within a population, and this time correlates with the time at which the ensuing spore germinates and in its ability to resume growth in various nutrients [9]. Spores germinate in reponse to low concentrations of nutrients at variable times [7], and this variability is partially explained by the number of germinant receptors in each spore [10]. Even in the absence of germinants, a small fraction of spores germinates spontaneously [11, 12]. These phenomena of phenotypic heterogeneity have been linked to variations in gene expression [8, 9, 12, 13]. However, the remarkable dynamics of metabolism during sporulation and germination have not been measured in individual cells, due to lack of a suitable technique.

The dynamics of energy metabolism can be measured in single vegetative bacteria using several *in vivo* methods, most of which are based on fluorescence [14]. These are difficult to apply to spores due to their strong autofluorescence, primarily from calcium dipicolinate, NADH, and flavins [15, 16]. The intracellular redox potential drives all pathways of energy metabolism other than fermentation [17, 18], and has been measured *in vivo* in single bacterial cells using a custom-fabricated microelectrode coupled to an optical trap [19]. However, this technique requires a specialized device, operates in anaerobic conditions, and only measures a small number of cells.

An alternative approach uses the bacterial luminescence system [20]. An enzyme, luciferase, encoded by the *luxAB* genes, catalyzes the light-emitting oxidation of a substrate, luciferin, which can be supplied exogenously or synthesized endogenously by expression of the *luxCDE* genes. The oxidation reaction’s rate depends on the reduced form of the electron transporter flavin mononucleotide (FMNH_2_), whose concentration is determined by the intracellular reducing power [21]. Thus, the rate of light production directly relates to the redox potential. In addition, since the emission of light occurs without external illumination, bioluminescent techniques can be considered “zero background”, and therefore offer improved sensitivity as compared to fluorescent techniques. Here we describe a microscope designed to capture the dynamics of bioluminescence emitted by single cells, which we used to non-invasively assay metabolic activity of single germinating spores of *Bacillus subtilis*.

## Results

### Bioluminescence microscopy of single spores

To date, the bioluminescence dynamics of sporulating and germinating cells have been measured only in populations, typically using a photomultiplier tube for photon counting [22, 23]. A *B. subtilis* strain that expresses the *lux* genes under a sporulation-specific promoter (P_*sspB*_) produces spores that contain a sufficient number of luciferase enzymes to emit detectable light only upon germination [23]. Prior to germination, spores do not emit detectable light, presumably due to their dormant metabolic state.

The feasibility of measuring bioluminescence dynamics in single germinating spores is determined by the rate of photon emission, which must be sufficiently high to be resolved over measurement noise. This rate increases with higher expression of the *luxABCDE* genes, as has recently been achieved in *B. subtilis* by modifying the ribosomal binding sites upstream of each gene [24]. The rate of bioluminescent photon emission at the population level has been reported in relative light units, or RLUs [24, 25], which are defined differently for various devices. RLUs can be converted to units of photons per time per cell by calibration with a luminescent standard [26], although this calibration has not been included in the published results. As a rough guide, microscopy of individual metabolically active *Escherichia coli, Synechoccocus elongatus, Vibrio fischeri*, and *Vibrio harveyi* cells expressing the lux system typically measures ∼60 photons min^-1^ cell^-1^ [27-29]. Our preliminary analysis suggested that a microscope utilizing recent improvements in CCD technology could resolve bioluminescence signals as dim as 1 photons min^-1^ spore^-1^ (SI Appendix).

We therefore constructed a custom microscope for measuring the dynamics of bioluminescence in single spores (Fig. 1A, Materials and Methods). The microscope includes an optical path for bright field imaging, which is used to localize spores and measure their refractility, as described below. The setup consists of an inverted microscope with a high-NA 63X objective and a cooled, back-illuminated CCD camera. We used 10 min exposure times, sufficient to resolve germination dynamics while achieving sensitivity to permit measurements from single germinating spores. Over a period of 10 min, Brownian motion displaces a spore by a root-mean-square displacement of approximately 50 μm. To prevent such displacements, we trapped the spores by using hydrophobic glass coverslips. With this method, the displacement of spores over 10 min periods was below the optical resolution of the bright field images (<160 nm). The sample was held in a perfusion chamber, which dynamically controlled the chemical environment (Fig. 1B), and the temperature at the sample was controlled to ±0.05 °C by an air-to-air heat exchanger.

We initially investigated the dynamics of bioluminescence emitted by single *B. subtilis* spores as they germinated in response to L-alanine. These spores (JDB3780) expressed the *luxABCDE* genes under the control of a sporulation-specific promoter. A field of approximately 900 spores was imaged in bright field and luminescence every 10 min for 1.5 h (Fig. 1C). Prior to the induction of germination, the dormant spores appeared dark, with high contrast, and produced no resolvable luminescence signal (Figs. 1D and E). L-alanine was added to the perfusion media at a concentration of 100 μM, and the ensuing germination response was recorded for 15 h.

**Fig. 1.**
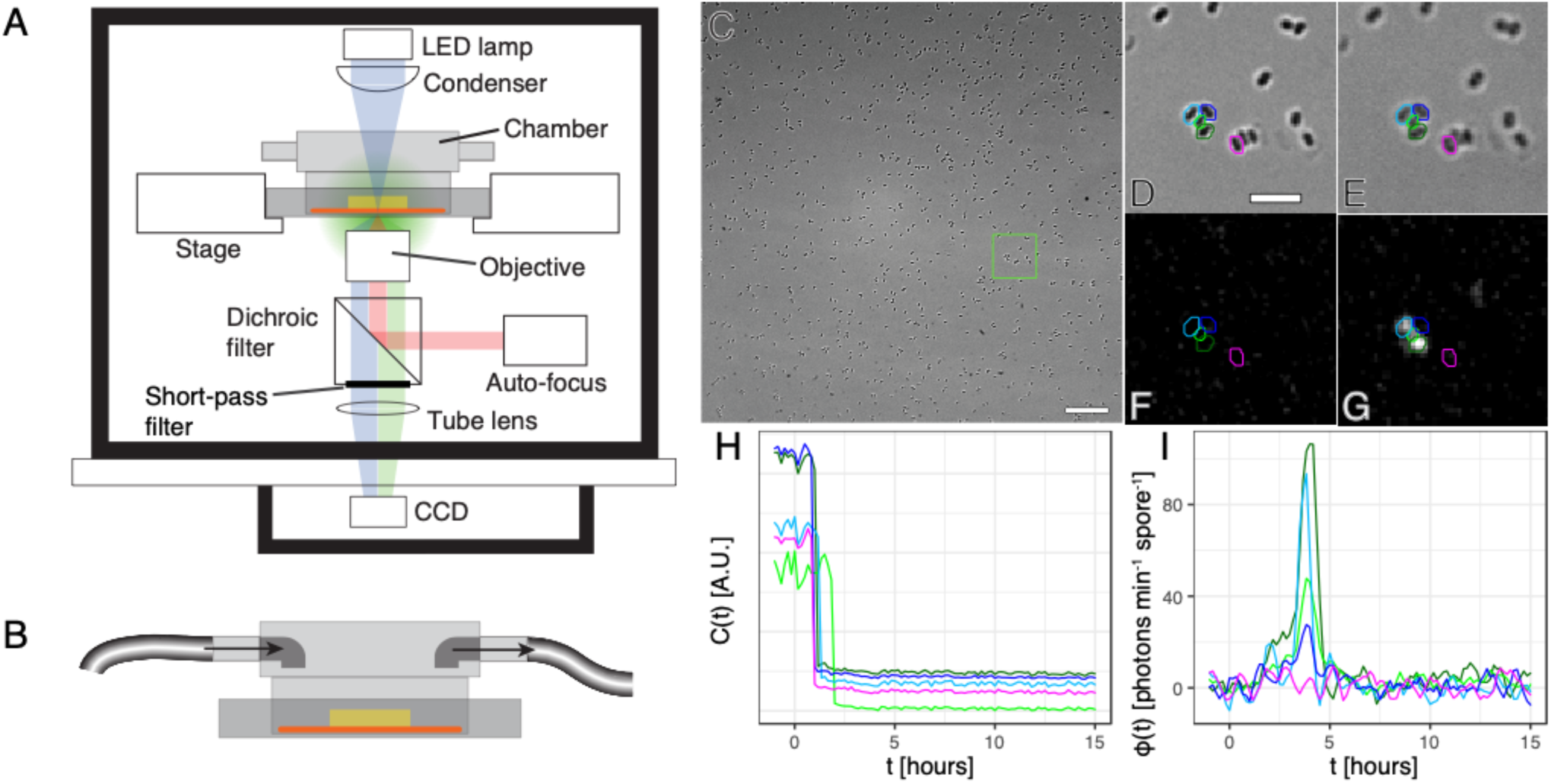
Schematic of bioluminescence microscope and perfusion chamber with example images and time series. **(A)** Microscope schematic showing 3 light paths: bright field (blue), luminescence (green), and autofocus infrared (red). For bright field imaging (blue path), the external illumination is focused near the bottom of the agarose pad (yellow slab), where sample cells are sandwiched against a glass coverslip. The sample emits luminescence (green path) in all directions. A temperature-controlled, light-tight box (thick black lines) encloses the microscope. **(B)** The perfusion chamber flows medium around the agarose pad (yellow) that traps sample cells against a glass coverslip on the chamber bottom, sealed by an O-ring (orange). **(C)** Bright field image of P_*sspB*_-*luxABCDE* spores, before the addition of germinant (time 0 h). Highly refractile dormant spores appear dark, *i.e.*, high contrast. The field of view contains approximately 900 spores. Scale bar: 20 μm. **(D**,**E)** The subregion bounded by the green box is shown in panels (D) (bright field) and (E) (luminescence), showing the lack of luminescence signal from dormant spores. **(F**,**G)** The same region is depicted 3.5 h after the addition of 100 μM L-alanine, showing spores’ typical loss of bright field contrast upon germination (F), and the corresponding increase in luminescence signal (G) that occurs in some cells after germination. **(H**,**I)** Time series of bright field contrast (H) and luminescent photon counts (I) of the 5 representative spores outlined in (D)-(G). The rapid loss of contrast shown in panel (H) defines the time of germination, *τ*_*g*_, for each spore. Colored cell outlines are dilated by 1 pixel for visualization. Panel (D) scale bar: 5 μm.

After an initial lag following the addition of L-alanine, most spores showed a significant decrease of contrast in bright field, and many produced luminescence well above the noise background (Figs. 1F and G). The predominant source of measurement noise was sensor read noise; we found that its contribution could be reduced significantly by applying a temporal low-pass filter with a cutoff period of 36 min (SI Appendix). Sensor read noise has a Gaussian distribution, and can therefore produce estimated photon counts that are negative [30]. We determined that the technique had a sensitivity limit of 0.4 photons min^-1^ spore^-1^ (SI Appendix).

The dynamics of individual spores in the bright field and luminescence images were analyzed, resulting in time series of bright field contrast and estimated luminescent photon counts for individual spores (Figs. 1H and I). Rapid loss of contrast after the addition of L-alanine was evident in most spores, indicating an early event in germination, namely, the release of calcium dipicolinate from the spore [31]. Spores differed significantly in their time of germination (Fig. 1H) and in their bioluminescence signals (Fig. 1I).

### Bioluminescence dynamics are synchronized with germination time

In order to compare the measurements in the microscope to an established method, we used a 96-well plate reader to measure the population-averaged bioluminescence dynamics of spores germinating in response to the addition of L-alanine at 5 different concentrations (1 μM, 10 μM, 100 μM, 1 mM, and 10 mM) (Fig. 2A). Time-lapse experiments were performed in the microscope setup at the same L-alanine concentrations, and the population-averaged bioluminescence dynamics were calculated (Fig. 2B). The dynamics as measured by both methods shared some features: Signal intensity generally increased with increasing L-alanine concentration, from total darkness at 1 μM, to a peak level at 1 mM, then decreased slightly at 10 mM (Fig. 2C). The L-alanine concentration also affected the temporal patterns of the dynamics: In the measurements performed with the microscope, a peak in the bioluminescence signal was evident approximately 1 h after germination with 10 mM, 1 mM, and 100 μM L-alanine, but this peak was not observed following germination with 10 μM L-alanine (Fig. 2B). Although much less pronounced, this early peak was also apparent in the plate reader measurements after germination with 10 mM and 1 mM L-alanine; with 100 μM and 10 μM L-alanine, no peak was resolved, but a “shoulder” at 40 min was apparent in both conditions (Fig. 2A). The primary difference in the dynamics measured by the two methods was the much higher intensity of the early peak measured in the microscope. We observed similar differences caused by varying the spore density in the plate reader experiments (Fig. S1), which are likely due to chemical interactions between germinating spores. This effect can explain the differences observed between the dynamics measured in the plate reader, at a density of 3×10^7^ spores ml^-1^, and in the microscope, at a density of less than 10^5^ spores ml^-1^.

**Fig. 2.**
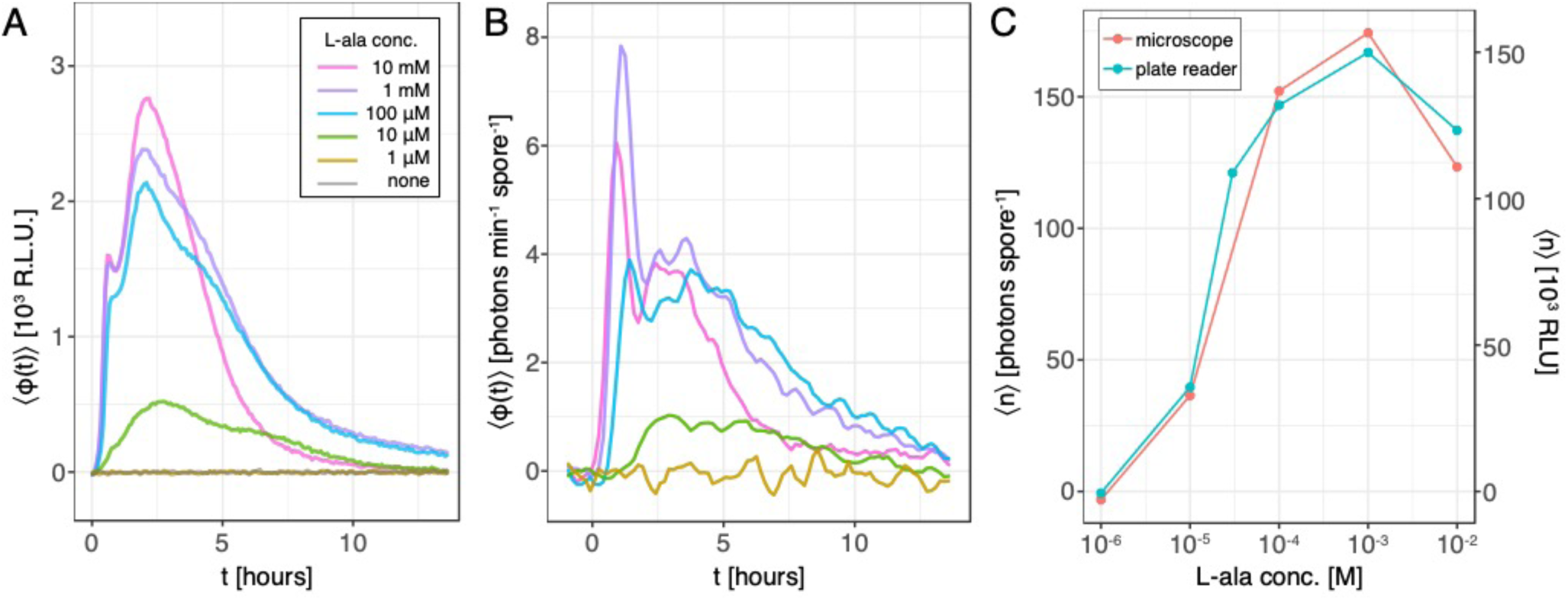
Bioluminescence dynamics of spore populations germinating in response to L-alanine. **(A)** The bioluminescence emitted by P_*sspB*_-*luxABCDE* spores germinating in a range of L-alanine concentrations was assayed in a 96-well plate reader. Each well contained approximately 6×10^6^ spores. Lines depict the mean signal over 3 replicates; the standard deviation was negligible on this scale. **(B)** The bioluminescence signal *ϕ*(*t*) was measured in individual spores germinating in response to various concentrations of L-alanine added at time 0 h. The mean signal over all spores (>600 individuals per concentration) is plotted at each time point. **(C)** The bioluminescence signals in (A) and (B) were integrated over the period between 0 and 12 h after the addition of alanine to calculate the population-averaged photon counts *n*, plotted as a function of alanine concentration.

The observed population-averaged bioluminescence dynamics arise from temporal fluctuations in individual spores as well as variability in the population, and these two sources can be separated using the single-spore technique. In particular, individual spores germinate at different times, with large variability observed at low L-alanine concentrations [32]. With automated image analysis, we identified the time *τ*_g_ of rapid contrast loss in each germinating spore (SI Appendix), and interpreted *τ*_g_ as the time at which each spore started germination. We considered the statistics of germination dynamics by calculating the fraction of the population that had not yet germinated by time *t*: This is the tail distribution function of germination time, P(*τ*_g_>*t*) (Fig. 3A). At L-alanine concentrations of 100 μM and higher, most spores germinated within 1 h of induction, and over 99% of spores germinated within 13 h. In contrast, spores germinated much more slowly in response to the addition of 10 μM L-alanine, with only half of all spores germinating within 13 h. For the 1 μM L-alanine concentration, we did not observe any germination for spores up to 82 h after induction.

**Fig. 3.**
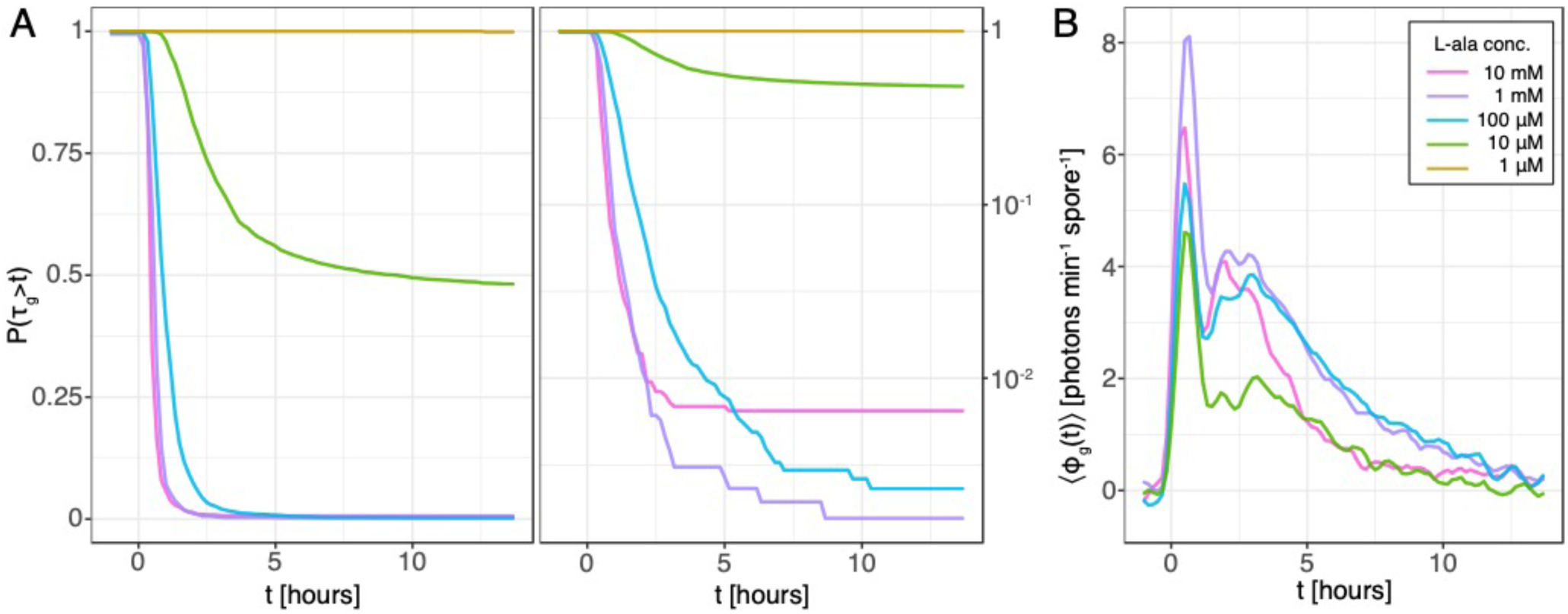
Time of germination varies in populations and synchronizes with bioluminescence dynamics. Dormant spores were exposed to 5 concentrations of L-alanine at time 0 h. **(A)** The fraction of spores that had not yet germinated by time *t*, P(*τ*_g_>*t*), is plotted on a linear scale (left) and a log scale (right). An offset is added to avoid divergence of zero values on the log scale. **(B)** The luminescence signal of each spore as a function of time elapsed since the time of germination is *ϕ*_g_(*t*) = *ϕ*(*t*-*τ*_g_), defined for spores that germinated during the acquisition interval. The population average of *ϕ*_g_(*t*) is plotted for each concentration of L-alanine. Note that no spores germinated in 1 μM L-alanine and so no curve for that condition appears in (B).

The distributions of germination time *τ*_g_ reveal significant variability in the population of spores, especially evident in the 10 μM L-alanine condition. To investigate the relationship between this variability and the bioluminescence dynamics, we considered the bioluminescence of each cell as a function of time elapsed since germination: *ϕ*_g_(*t*) = *ϕ*(*t*-*τ*_g_), defined for all spores that germinated during the observation interval. Note that the germination time *τ*_g_ was determined exclusively by the bright field images, through a procedure that was independent of the bioluminescence dynamics (SI Appendix). The population averages of *ϕ*_g_(*t*) in each germination condition indicate significant synchronization of the bioluminescence dynamics with the time of germination, as evidenced by the qualitatively similar temporal patterns in the 4 conditions for which germination was observed (Fig. 3B). In particular, we observe remarkable alignment in the timing of a primary peak at *t* ≈ 40 min, the appearance of a secondary peak between *t* = 2 h and *t* = 3 h, and similar profiles of signal attenuation at later times, possibly due to degradation of the lux enzymes (Fig. S2).

### Bioluminescence varies significantly across spores

Aligning the dynamics to the time of germination *τ*_g_ accounted for the occurrence and timing of germination as a source of the variability observed in the bioluminescence dynamics. Significant variability remained in the aligned dynamics *ϕ*_g_(*t*) (Fig. 4A). We analyzed the variability of *ϕ*_g_(*t*) over spores to answer two questions: First, what are the temporal correlations of the bioluminescence dynamics in individual spores? Second, how much of the variability in bioluminescence is due to variations in the factors that affect the reaction chemistry, namely, the redox potential, luciferin concentration, and luciferase copy number?

**Fig. 4.**
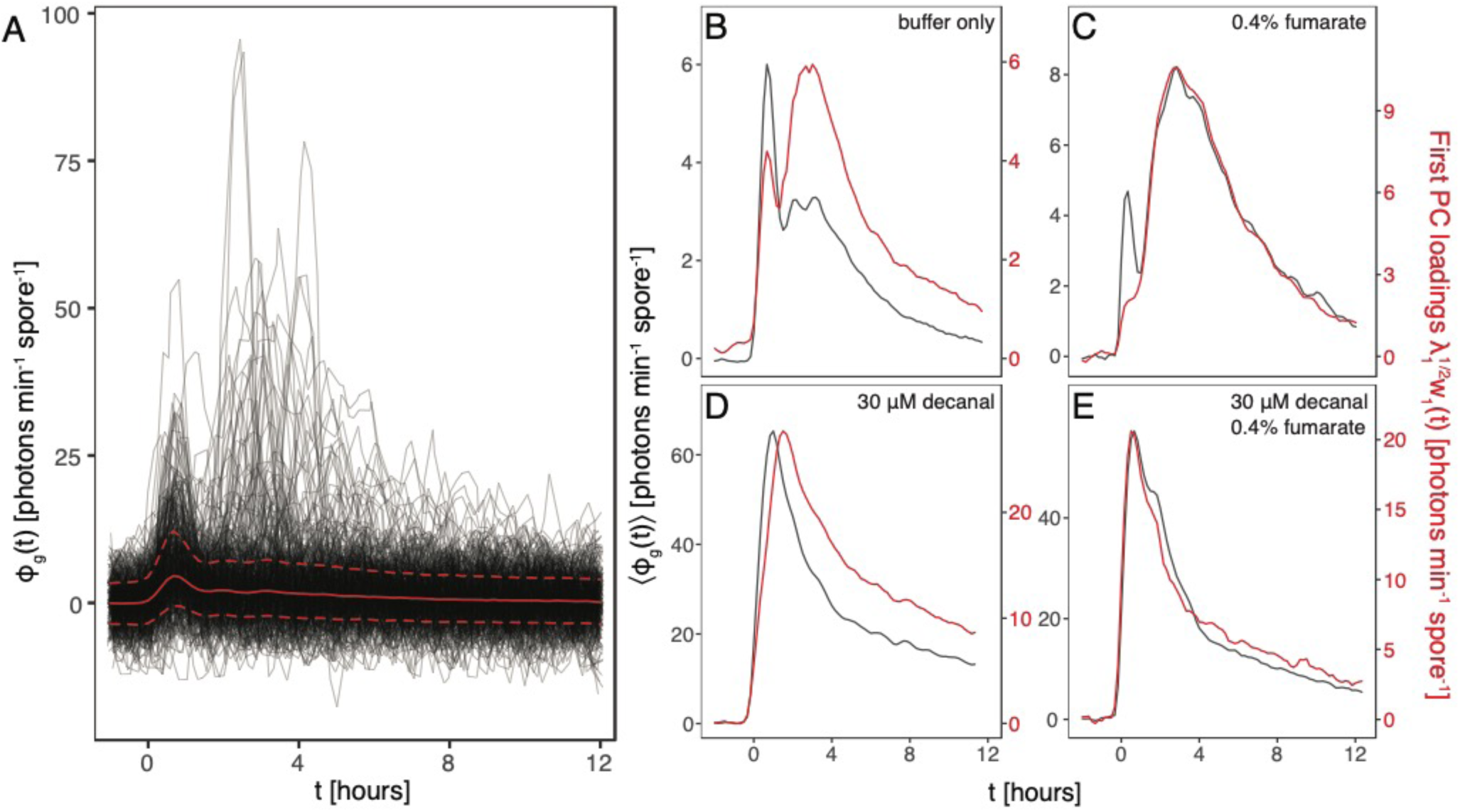
Variability in bioluminescence dynamics over spores. **(A)** Bioluminescence as a function of time since germination is analyzed in terms of variability over individuals. Data include all spores that germinated in response to L-alanine at 4 concentrations (Table 1). The time series of *ϕ*_g_(*t*) for a randomly chosen subset of 500 cells are plotted as thin black lines. The solid red line indicates the median over the 11,000 cells comprising the data set, and the dashed red lines indicate the 16% and 84% quartiles. (**B-E**) Principal component analysis was performed on time series *ϕ*_g_(*t*) from 4 experimental conditions: germination buffer only (B), 0.4% sodium fumarate (C), 30 μM decanal (D), and 30 μM decanal plus 0.4% sodium fumarate (E). The dark gray line shows the population average ⟨*ϕ*_g_(*t*)⟩ for each condition. The first principal component for each condition explained 30%, 55%, 68%, and 58% of the total variance, respectively. The red line shows the loadings of the first principle component, *i.e., λ*_1_^1/2^*w*_1_(*t*), where *λ*_1_ is the largest eigenvalue of the covariance matrix, and *w*_1_ is the first principle component, scaled to unit norm. The average and loadings are plotted on separate axes. The first principal component and the population average were highly correlated in all conditions: defining *r*=cor(*w*_1_(*t*),⟨*ϕ*_g_(*t*)⟩), *r*_buffer-only_=84%, *r*_fumarate_=97%, *r*_decanal_=89%, *r*_decanal+fumarate_=99% (all *p*-values <2.2×10^-16^).

To analyze temporal correlations in the bioluminescence dynamics, we calculated the temporal covariance matrix *C*(*s,t*) from the photon counts *ϕ*_g_ at times *s* and *t* (SI Appendix). This calculation was performed over all germinating spores from the 4 concentrations of L-alanine for which germination was observed. The covariance matrix was analyzed using principal component analysis (PCA): PCA applied to time series in this manner is often referred to as empirical orthogonal function analysis [33]. The first principal component explained 30% of the variance over individuals, while additional components each explained only a few percent of the total variance (Fig. S3). Therefore, we focused on the pattern of correlations described by the first component, which had temporal structure similar to the population-averaged dynamics (Fig. 4B). This similarity indicates that the dynamics in most spores were well-described by the mean dynamics multiplied by a coefficient specific to each spore, which we call its amplitude. This simple mode of variation was useful for quantifying the contributions from different sources, as described below.

**Table 1.**
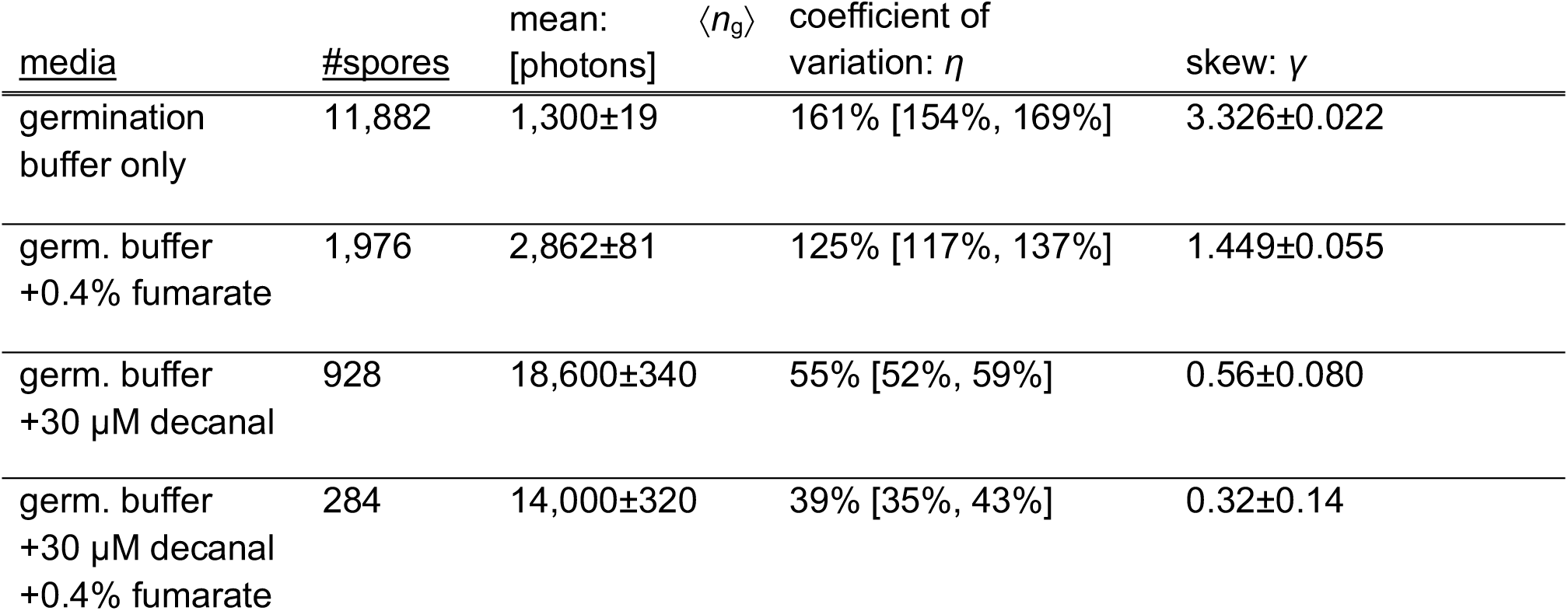
Bioluminescence signal amplitude statistics. Columns denote media, the number of spores analyzed, and the statistics of *n*_g_, the photon counts per spore integrated over the 12 h after germination. The ± signs indicate 1 standard error. The bracketed interval in the coefficient of variation column is the 95% confidence interval.

### Bioluminescence variability arises from variations in redox potential and luciferin concentration

Besides the timing of germination, several factors could contribute to variability in the bioluminescence dynamics. The redox potential and concentration of luciferin in each cell are two obvious candidates. To investigate these, we performed an experiment with fumarate added to the germination buffer, in order to drive the citric acid cycle and thus increase the intracellular redox potential. In a second experiment, decanal was added as an exogenous luciferin. In a third experiment, both fumarate and decanal were added to the germination buffer. Spores germinated in response to the addition of 1 mM L-alanine. We calculated the population-averaged bioluminescence dynamics and performed PCA for each data set independently (Figs. 4C-E).

In the presence of fumarate, the population-averaged bioluminescence dynamics ⟨*ϕ*_g_(*t*)⟩ exhibited 2 distinct peaks, approximately 40 min and 3 h after germination (Fig. 4C). These dynamics appear to be similar to those observed in germination buffer without fumarate (Fig. 4B), with two major differences: The addition of fumarate increased the intensity of the second peak, and strongly attenuated the spore-to-spore variability during the first peak, as indicated by the disappearance of the first peak from the first principal component. The addition of exogenous luciferin (decanal), both with and without fumarate, resulted in population-averaged bioluminescence dynamics with a single peak, occurring approximately 40 min after germination, with intensity approximately 10-fold higher than the initial peak observed in the absence of exogenous luciferin (Figs. 4D and 4E).

In all conditions, PCA revealed a single significant component of variation (Fig. S3), and in each condition, this mode’s eigenvector was strongly correlated with the average dynamics (Figs. 4B-E). Thus the primary mode of variation in all conditions was variability in amplitude. This simple pattern facilitates the quantitation of variability, since the contribution of each spore’s signal to the total variability can be summarized by a single variable: the signal amplitude. Thus we calculated for each germinating spore the total number of photons counted during the 12 h following germination, *n*_g_.

The mean value of *n*_g_ varied significantly in different conditions. Therefore, to compare variability in *n*_g_ across conditions, we considered the ratio of the standard deviation to the mean, 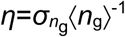, also known as the coefficient of variation. We also considered the asymmetry in the distribution of *n*_g_ by calculating its skew, *γ*. The estimated statistics are summarized in Table 1. Variability was high for spores germinating in the absence of exogenous factors (*η*=160%), and the distribution of *n*_g_ had strong positive skew (*γ*=3.3). The addition of fumarate significantly decreased the coefficient of variation and the skew (*η*=125%, *γ*=1.4). The addition of decanal reduced the variability and skew more strongly (*η*=55%, *γ*=0.6). The addition of both fumarate and decanal had the largest effect (*η*=39%, *γ*=0.32). We conclude that the large amount of variability in bioluminescence signal observed in spores germinating in the absence of exogenous factors was predominantly due to variability in both intracellular luciferin concentration and redox potential.

The bioluminescence dynamics following germination are influenced by metabolic changes that occur early in germination, including the initiation of electron transport [23, 34]. In addition to inducing germination, L-alanine may also fuel energy metabolism [9]. To investigate this possibility, we considered the photon counts *n*_g_ from spores germinating in response to 4 concentrations of L-alanine, in germination buffer (Fig. S4). The population-average ⟨*n*_g_⟩ increased 2-fold as the L-alanine concentration increased from 10 μM to 1 mM, then decreased at 10 mM. The variation in amplitude with L-alanine concentration arose primarily from a subpopulation of the brightest spores (Fig. S5). The weak, non-monotonic dependence of the bioluminescence signal on L-alanine concentration indicates that exogenous L-alanine does not significantly drive energy metabolism in germinating spores.

## Discussion

The spore is an extreme phenotypic state that allows an organism to survive extended periods of starvation, drought, and radiation. Metabolism decreases to extremely low levels during sporulation, then rapidly increases during germination. Laboratory studies of populations undergoing the sporulation and germination transitions have also revealed large amounts of variability over individuals, suggesting that phenotypic diversification in natural spore-forming populations is a bet-hedging strategy for surviving in fluctuating environments. It is therefore important to study metabolic dynamics in individual spores.

No previously described methods for quantifying metabolic dynamics in single cells are applicable to *Bacillus* spores. Therefore, we developed a microscopic technique for this purpose, using a bioluminescent reporter for the intracellular redox potential. In this paper, we measured bioluminescence dynamics from individual spores germinating in response to L-alanine, which is both a germinant and a potential nutrient. Comparing the bioluminescence dynamics measured in individuals to the population average, we exposed significant variability in the population. A major source of variability was shown to be associated with differences in the timing of germination. We could remove this contribution from the bioluminescence dynamics by simultaneously using bright field microscopy to precisely determine the time of germination of each spore. The bioluminescence dynamics could then be analyzed for its temporal correlations. The analysis indicated that the primary mode of variation was associated with the amplitude of the bioluminescence signal produced by each spore.

By performing experiments at different concentrations of L-alanine, we were able to show that it was not a major source of variations in bioluminescence amplitude, suggesting that metabolism in germinating spores was driven not by L-alanine, but rather by endogenous energy sources stored during sporulation. Through the addition of exogenous nutrients (fumarate) and bioluminescence substrate (luciferin), we were able to decompose the variability in bioluminescence amplitude into two main sources: the intracellular redox potential and luciferin concentration. Both of these sources relate to energy metabolism, as the redox potential drives ATP synthesis, and the biosynthesis of luciferin depends on both ATP and NADPH [35]. Our results show that we can begin to understand the metabolic transitions between the growing and dormant states at the level of single cells.

We foresee two important extensions of the current work. First, one can study the variability of metabolic dynamics in fluctuating environments. Vegetative cells can be tracked as they sporulate in response to experimentally imposed starvation, and later germinate and outgrow in response to nutrients [9]. By simultaneously measuring the bioluminescence dynamics in this type of experiment, we can study the correlations between spores’ timing of sporulation, timing of germination, ability to outgrow, and metabolic state. Environments also fluctuate due to biotic interactions. Bacterial spores contain high concentrations of calcium dipicolinate, a compound that affects resistance and dormancy, but also induces germination in dormant spores [36]. Because spores release large amounts of calcium dipicolinate during germination [37], the local concentration of spores can affect their germination behavior. Such interactions may explain the differences we observed in the bioluminescence dynamics with respect to spore density (Figs. 2A and B, Fig. S1). Interactions between spores and their effects on metabolism could be studied systematically by analyzing the spatial statistics of germination timing and bioluminescence.

A second extension is to study the thermodynamic nature of dormant states. The maintenance of order in living matter seems to require energy expenditure, in order to counteract spontaneously occurring degradation [38, 39]. Therefore, a central question arises: What are the energetic limits for survival? This question has primarily been studied in microbial communities from low-energy environments, including deep sediments, permafrost, and subglacial lakes [40-42]. Metabolic activity in such environments, averaged over the community, has been found to be reduced by 5 to 6 orders of magnitude compared to the state of growth when nutrients are supplied [40, 41]. These low metabolic rates are approximately equal to the energetic costs of repairing molecular damage to proteins and DNA at the corresponding temperatures [41, 42]. However, these analyses included contributions from cells with metabolic activity that was potentially significantly lower than the population average, including dormant spores [43]. Therefore, spores likely represent the most extreme case of metabolic dormancy in nature.

Most studies of metabolism in dormant spores have not detected activity [44-47]. Extraodinarily sensitive measurements of metabolism in populations of dormant spores using radiochemical labelling of metabolites did detect activity, reduced by over 5 orders of magnitude compared to the vegetative state [4]. However, the detected metabolism could have been due to a rare subpopulation of germinating spores [4, 11, 12], and therefore can serve only as an upper bound on the rate of metabolic activity in dormant spores. By resolving individual spontaneous germination events, bioluminescence microscopy could achieve unprecedented sensitivity. The intensity of bioluminescence is linear in the concentration of FMNH_2_ at low concentrations [48], so its utility as a metabolic reporter should extend to cells with very low metabolic rates, such as dormant spores. A measurement on a single spore at a single time point with signal-to-noise ratio *ξ* can be averaged over *N* individuals and *T* time points to produce a bulk measurement with signal-to-noise ratio *ξ*(*NT*)^1/2^. For our method applied to germinating spores, *ξ* is ∼20 (SI Appendix). This suggests that bioluminescence measurements could achieve sensitivity concomitant with a 10^5^-fold reduction in signal compared to germinating spores by measuring, for example, a population of 10^4^ dormant spores over an interval of 18 d, which is feasible using the setup described here.

A possible mechanism for long-term survival without any metabolic activity is the formation of a thermodynamically metastable state. The glassy state is an example of metastability that has been invoked to explain spore dormancy [49, 50], as macromolecules in a glass change conformation extremely slowly [51]. Therefore, a glassy environment could in principle reduce the rate of molecular damage to proteins and DNA, while also reducing the activity of metabolic enzymes and thus causing dormancy. Direct characterization of glassiness in spores is difficult, as the internal structure of the spore is highly heterogeneous [52]. Instead, one can study a feature of spores from the perspective of metastability, namely, the statistics of spontaneous germination and its dependence on environmental conditions such as temperature. This approach should be possible with our technique, which would also quantify possible differences in metabolism between spores germinating spontaneously and those germinating in response to nutrients.

## Acknowledgments

We thank members of the Dworkin and S. Leibler labs for helpful discussions. ZF was supported by a grant from Simons Foundation to S. Leibler through Rockefeller University Grant 345430 and JD was supported by NIH GM122146.

## Materials and Methods

### Strains

JDB3780 (168 *trpC2 sacA::*P_*sspB*_*-luxABCDE::cm*) was constructed by transforming genomic DNA from RL4921 (PY79 *sacA::*P_*sspB*_*-luxABCDE::cm*) into JDB1772 (168 *trpC2).*

### Sporulation protocol

Spores were prepared in Difco sporulation media (DSM). Single colonies grown on LB agar were used to inoculate 3 ml cultures in test tubes. Cultures were grown with shaking at 30 °C for 3 h. 30 μl aliquots were used to inoculate 30 ml cultures in baffled flasks. After growing with shaking at 30 °C for 36 h, spores were purified by spinning at 5,400 g for 10 min at 4 °C and resuspending in sterile water. Cold water washes were repeated 4 times, yielding samples that were predominantly refractile spores (>90%). Samples were stored at 4 °C.

### Plate reader assay

To assay bioluminescence dynamics in response to various concentrations of L-alanine, dormant spores (JDB3780) were suspended in 25 mM Tris-HCl buffer at pH 8.2, at a density of 3×10^7^ spores ml^-1^. 200 μl of spore suspension was pipetted into wells of a 96-microwell plate (Nunc 236108). In order to avoid both signal contamination from neighboring wells as well as the influence of thermal gradients, wells were filled only in the plate interior, in a checkerboard pattern. 2 μl of L-alanine solutions at concentrations between 1 M and 1 mM were added to the appropriate wells. Each germinant condition was prepared in 3 replicate wells. Three wells with 200 μl of buffer were included as blanks. The plate was sealed with a gas permeable, partially adhesive film (Uniscience C20613201), and positioned within the plate reader (Perkin Elmer Wallac Victor^2^ 1420-011). Luminescence measurements were made without an optical filter, normal aperture setting, and 10 s integration time. The temperature was maintained at 30 ± 1 °C. Between luminescence readings, the plate was shaken orbitally for 10 s with a 2 mm diameter at the slow speed setting. The plate reader assay of bioluminescence in the presence of chloramphenicol was performed using a similar procedure, with spores of JDB3785 suspended at a density of 1.2×10^8^ spores ml^-1^ and germinated with 1 mM L-alanine.

### Reagents and Media

All germination experiments were conducted in germination buffer (25 mM Tris-HCl, pH 8.2). Decanal was dissolved at a concentration of 15 mM in a 1:1 solution of methanol and water.

### Microscopy

#### Sample preparation

All sample preparation was conducted with sterile technique at the bench.

Adhesion of dormant and germinating spores to glass coverslips was improved by rendering the glass surface hydrophobic. Coverslips were treated with vapor deposition of hexamethyldisilazane (Alfa Aesar L16519AE) in a sealed chamber for 1 h, followed by baking at 80 °C for 1 h.

Agarose pads were prepared by pipetting 700 μl of molten 4% low-melt agarose (Fisher BP165-25) between two 22 mm diameter glass coverslips. After 1 h at room temperature, the agarose pad was exposed by removing the top coverslip, and dried for 5 min.

Prior to spotting the pad, dormant spores were washed twice with water (5400 g, 10 min, 4 °C). 2 μl of the spore suspension was pipetted onto the center of the dried pad, and the spot was allowed to completely evaporate (∼10 min). The pad was then separated from the bottom coverslip, and inverted onto a hydrophobic glass coverslip (30 mm diameter, #1.5 thickness, Bioptechs). The coverslip-bonded pad was left to evaporate for 30 min, then assembled into the bottom of the perfusion chamber (Bioptechs, ICD with heated lid, gas and media perfusion). Molten 4% low-melt agarose was added to the exposed portion of the coverslip, and allowed to gel for 30 min. Creating a contiguous layer of agarose in this manner was found to attenuate coverslip bounce resulting from the perfusion flow.

#### Perfusion

A peristaltic pump (Cole-Parmer, Masterflex L/S digital miniflex, dual-channel) was used to draw media from a reservoir bottle, through sterile tubing (Cole-Parmer, C-Flex #13, i.d. 0.8 mm), into the perfusion chamber, and finally into a waste reservoir bottle. Tubing was connected with standard Luer locks, and the bottle openings were wrapped tightly with Parafilm and covered in aluminum foil. At the beginning of each experiment and whenever the medium was changed (for example, when germinant was added), the flow was primed at 1 ml min^-1^. Otherwise, the flow rate was 50 μl min^-1^.

Prior to conducting each imaging experiment, the perfusion tubing was sterilized by flowing 100 ml 20% bleach, followed by 100 ml isopropanol, followed by 5 subsequent bottles containing 250 ml sterile, purified water (Milli-Q). The perfusion chamber was cleaned by rinsing with 20% bleach, isopropanol, and distilled water, then air dried.

#### Imaging

The imaging setup consisted of an inverted optical microscope (Zeiss Axiovert 135), a 63X 1.4 NA oil immersion DIC objective (Zeiss 1113-108) operated without the Wollaston prism, and a sensitive, back-illuminated, cooled CCD camera (Princeton Instruments, PIXIS 1024B).

A collimated, 445 nm LED (Thorlabs, Solis-445C) was used as external illumination for bright field imaging. For acquisition of bright field images, the LED was pulsed for 50 ms at low current (40 mA), and was otherwise fully off.

Autofocus was maintained via a pupil obscuration method applied to an infrared (780 nm) beam reflected from the glass-media interface (Applied Scientific Instrumentation, CRISP). The defocus detector controlled the focal position through a DC servo motor connected to the mechanical focusing shaft of the microscope (Applied Scientific Instrumentation, MFC-2000).

Luminescence images were acquired with 10 min exposure time. The readout was performed at 100 kHz (read noise 3.56 e^-^), high gain (1.00 e^-^ ADU^-1^), and with 2×2 binning. The CCD was cooled to −65 °C with a combination of thermoelectric and low-vibration liquid cooling (Princeton Instruments, CoolCUBE II). At this temperature, the dark current was measured at 10^-3^ e^-^ pixel^-1^ s^-1^.

The microscope and camera were enclosed inside a box made of styrofoam-lined black plastic. Exterior edges and corners of the box were covered with opaque black fabric and taped. A removable styrofoam-lined plastic panel provided access to the microscope and sample holder, and was held in place during data acquisition with Velcro straps and metal L-brackets. The temperature within the box was controlled by a thermoelectric device (TE Technology, AC-220), coupled to a PID feedback device (TE Technology, TC-720) monitoring a thermistor placed as close as possible to the perfusion chamber. The interior temperature of the box was maintained at 30±0.05 °C. The entire imaging apparatus was supported by a vibration-damped optical table, situated in a temperature-controlled room (28±0.1 °C).

### Image analysis and statistics

See SI Appendix.

## Supporting Materials and Methods

### Sensitivity limits of bioluminescence microscopy

Sensor read noise, dark current, and high energy particles such as cosmic rays generate a noise background for imaging. The signal measured from a steadily emitting source accumulates with increasing exposure time *t*. Noise due to dark current also accumulates with *t*, but modern cooled CCDs have very low dark current, so that sensor read noise dominates for *t* up to about 1 h. The number of high energy particle artifacts per image also increases with *t*. These artifacts can be identified automatically [1], so that pixels affected by these artifacts can be removed from subsequent analysis. From a practical perspective, *t* must be short enough to ensure that the majority of pixels are not affected by these artifacts. The flux of high energy particles from cosmic rays is about 1 cm^-2^ min^-1^ [1]. In our preliminary experiments, artifacts from high energy particles limited *t* to 10 min.

After removing artifacts due to high energy particles, imaging noise is dominated by sensor readout noise. We consider the following parameters: the average rate of photon emission per spore, *ψ*; the average number of pixels in the luminescence image of a spore, *N*; the RMS readout noise per pixel *σ*_r_; the exposure time *t*, and the collection efficiency of the imaging setup *s*. Note that the collection efficiency *s* is the ratio of the number of emitted photons to the number of electrons measured by the sensor. It includes geometric effects, *i.e.*, losses due to the collection geometry, reflections from optical surfaces, and the quantum yield of the sensor.

The signal to noise ratio for each pixel in the image of the spore, *χ*_p_, is the number of electrons per pixel per exposure arising from emitted luminescence, divided by the magnitude of the readout noise: *χ*_p_ = *sψtN*^-1^*σ*_r_^-1^. The signal from each spore is the sum over the signals from the pixels comprising its image. The readout noise is independent for each pixel, so the signal to noise ratio per spore, *χ*, equals *χ*_p_*N*^1/2^ = *sψtN*^-1/2^*σ*_r_^-1^. The sensitivity limit of the technique is taken to be the rate of photon emission per spore for which the signal to noise ratio *χ* is equal to 1: *ψ*_min_ = *s*^-1^*t*^-1^*N*^1/2^*σ*_r_.

Optimizing the sensitivity of the technique amounts to maximizing collection efficiency *s*, minimizing the number of pixels per spore *n*, and minimizing the sensor read noise *σ*_r_. Practically, *s* is less than 1/2 electrons per photon, since at least half of photons are emitted away from the collection objective. The numerical aperture of the objective should be as high as possible, to maximize collection of the remaining photons. The quantum efficiency of back-illuminated CCDs in the spectral range of bacterial bioluminescence (∼490 nm) typically exceeds 0.9 electrons per photon. Accounting for these effects, we estimated that we could achieve a value of *s* as high as 0.3 electrons per photon. For the camera we used in the microscope, the parameters were as follows: *s* = 0.3 e^-^ photon^-1^ (estimated), *t* = 10 min, *N* = 10 pixels, *σ*_r_ = 3.6 e^-^, which determines the sensitivity limit *ψ*_min_ ≈ 3.8 photons min^-1^. Published results typically report the number of measured photons per time (*γ*^-1^*sψ*, where *γ* is the gain, expressed in electrons per photon), rather than the number emitted per time (*ψ)*, and so we introduce a new symbol for the former rate: *Φ* = *γ*^-1^*sψ*, and estimate the sensitivity limit *Φ*_min_ = *γ*^-1^*sψ*_min_ ≈ 1 photons min^-1^ (*γ* = 1 e^-^ photon^-1^ for our setup).

We analyzed the signal to noise ratio of our measurements on a per-spore basis. The noise background was quantified by considering the statistics of luminescence signal from the reported experiments in regions of the image that did not contain spores. These regions were defined to each include the same number of pixels as a real spore. Twelve thousand such regions were analyzed, and the standard deviation of the estimated photon counts from each region was calculated: *σ* = 3.5 photons min^-1^. The bioluminescence dynamics per spore *Φ*(*t*) varied over time and across spores. Detection of the signal from a particular spore is primarily determined by that spore’s maximal signal. For the purposes of determining a single signal to noise ratio that would describe the typical sensitivity of the technique, we calculated the population-average of the temporal maximum bioluminescence signal:

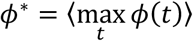

The signal to noise ratio is given by *χ* = *Φ*^*^*σ*^-1^. Its values were *χ* = 4.2 (germination buffer only), 5.1 (0.4% fumarate), 22 (30 μM decanal), and 18 (30 μM decanal + 0.4% fumarate).

### Image analysis

#### Background subtraction

Spatial inhomogeneities were observed in long exposure images, even those acquired in complete darkness without a luminescent sample. These inhomogeneities were found to be nearly constant over time and were treated as a form of background. To remove this background, a sequence of at least 20 images was acquired in darkness prior to each experiment, and the temporal median of this sequence, *i.e.*, the estimated background, was subtracted from each luminescence image subsequently acquired during the experiment.

#### Removal of artifacts due to high energy particles

Each luminescence image included several small regions of extremely bright pixels, due to high energy particles, such as cosmic rays, ionizing the CCD. In order to prevent these artifacts from corrupting the luminescence signal, we developed a routine to automatically detect pixels that appeared to be affected by such events. Pixels affected by artifacts had a particularly identifying feature: extreme brightness, that appeared rapidly in time and with very high spatial contrast. Temporal and spatial high-pass filters were applied to the images, and the resulting images were segmented, using a threshold chosen to separate the vast majority (>99%) of affected pixels from unaffected pixels. The affected regions were grown by 2 pixels to avoid any remaining signal contamination. The mean number of artifacts was 2.0 min^-1^, and the mean number of pixels affected by each artifact was 28.

#### Spore identification and segmentation

Spores were identified by determining the maximally stable extremal regions of the bright field image, using the VLFeat software library [2, 3]. The parameters of the method were tuned manually to identify >99% of spores, with a negligible false identification rate (<0.1%).

#### Determination of Germination Time *τ*_g_

The bright field contrast of each spore was determined by calculating the spatial gradient of each bright field image, ∇ *I*_BF_. The magnitude of the gradient was averaged over the pixels *p* corresponding to each spore to calculate its contrast:

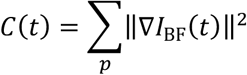

The dynamics of contrast of germinating spores typically appeared as noisy step function, declining from a high value prior to germination to a low value after. The time of germination *τ*_g_ was determined for each spore by fitting the measured contrast with a regression tree having a single split, using the rpart package for R, v. 4.1-15. The automatically determined *τ*_g_ was found to exactly match the value determined by manually considering the time-lapse images in 100 spores selected randomly from 4 data sets.

### Statistical analysis

#### Temporal filtering

Camera read noise and photon counting noise (shot noise) corrupt the counts of bioluminescence photons. Each of the noise sources is expected to have zero temporal autocorrelation, while the bioluminescence dynamics are expected to have significant autocorrelation. Defining the photon counts within each pixel *p* as *X*_*p*_(*t*), we expect the power spectral density of *X*_*p*_ to be different for pixels located within the image of a spore vs. pixels located outside (background). To text this, we estimated the power spectral density of *X*_*p*_(*t*) over 110,000 pixels within the image of spores (“signal”: *P*_S_(*ω*)) and over 200,000 pixels not within the image of spores (background: *P*_BG_(*ω*)). The power spectral density of the background was flat, while that of the signal was flat above *ω* ≈ 0.84 h^-1^. In order to filter the background noise without significantly blurring the signal dynamics, we applied a low-pass filter (sinc filter), with a cutoff period of 36 min, to the bioluminescence signal measured from each pixel. For each spore, the sum of these counts over all pixels inside its bright field image was calculated for each time point, resulting in *Φ*(*t*).

#### Principal component analysis

The covariance matrix *C*(*s,t*) was calculated over all germinating spores as:

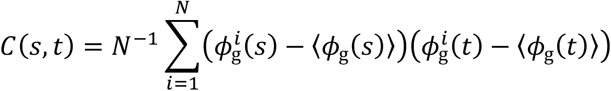

where 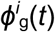 is the estimated rate of photons counted from spore *i* at time *t*, and *N* is the number of spores in the data set. The times *s* and *t* took values corresponding to 87 time points: 12 before germination (2 h), and 75 following germination (12.3 h). Principal component analysis was performed on the covariance matrix *C*(*s,t*), resulting in the eigenvalues *λ*_*i*_ and the eigenvectors *w*_*i*_(*t*). The eigenvalues have units (photons min^-1^)^2^ and the eigenvectors are unitless and have norm 1:

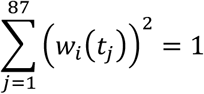

The temporal pattern of variation explained by mode *i* is quantified by *λ*_*i*_^*1/2*^*w*_*i*_(*t*), as is plotted in Figs. 4B-E for *i* = 1.

#### Statistics of *n*_g_

The number of photons counted per spore was calculated as

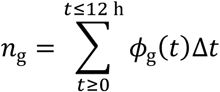

where Δ*t* = 10 min is the exposure time.

For Table 1, the estimates of the skew of *n*_g_ and its standard error were calculated for each data set using method 3 of Joanes & Gill [4]. The confidence interval of the coefficient of variation was estimated using the formula of McKay [5].

**Figure S1.**
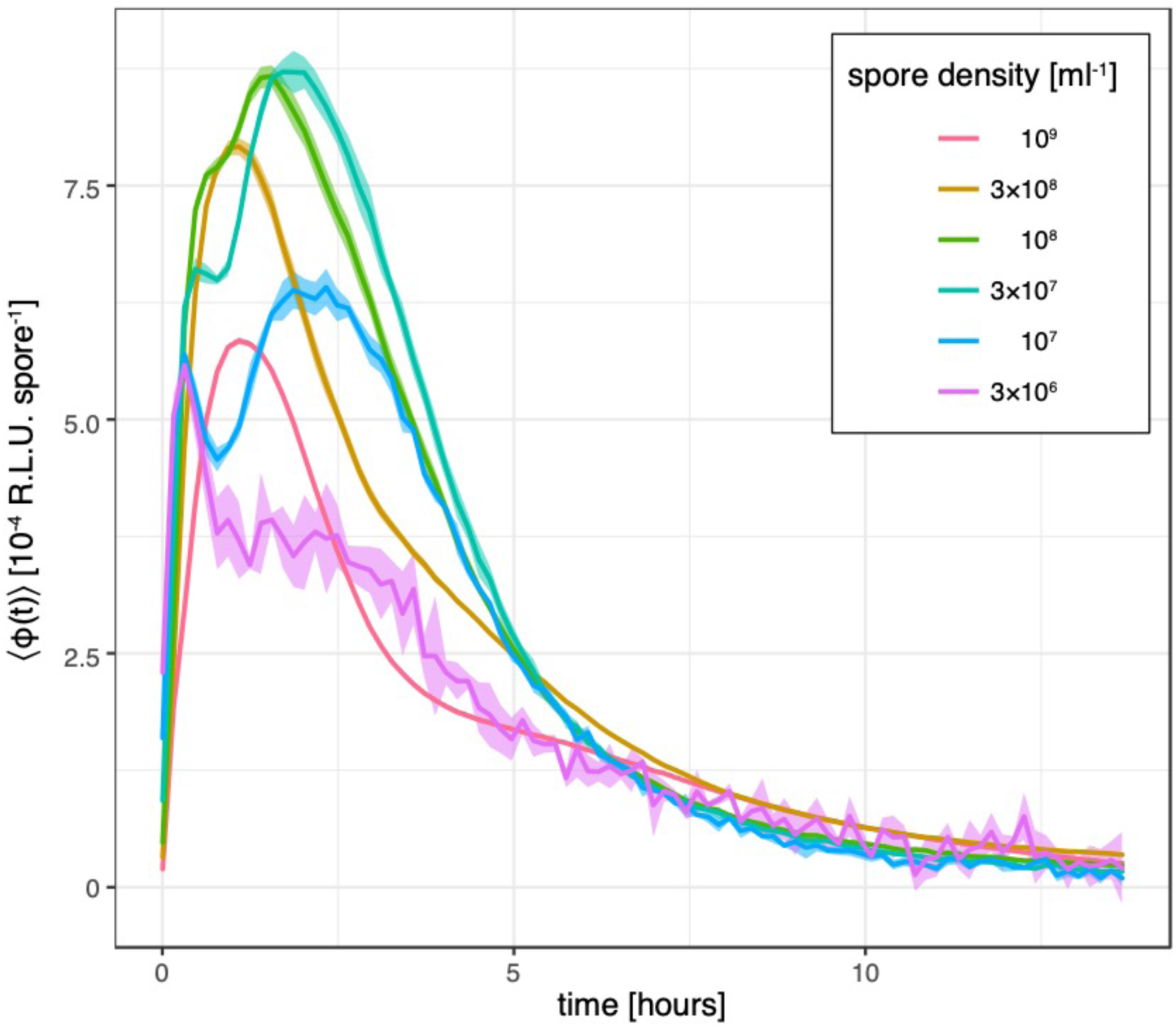
Bioluminescence dynamics depend on spore concentration. The population-averaged bioluminescence signal ⟨*ϕ*(*t*)⟩ was measured in a 96-well plate reader, at a range of spore densities, and normalized by the number of spores per well to facilitate comparison between densities. Spores were induced to germinate with 1 mM L-alanine. Lines depict the mean signal over 3 replicates and the shaded region encloses ±1 standard deviation. Spore density in the experiments performed with the microscope was less than 10^5^ ml^-1^.

**Figure S2.**
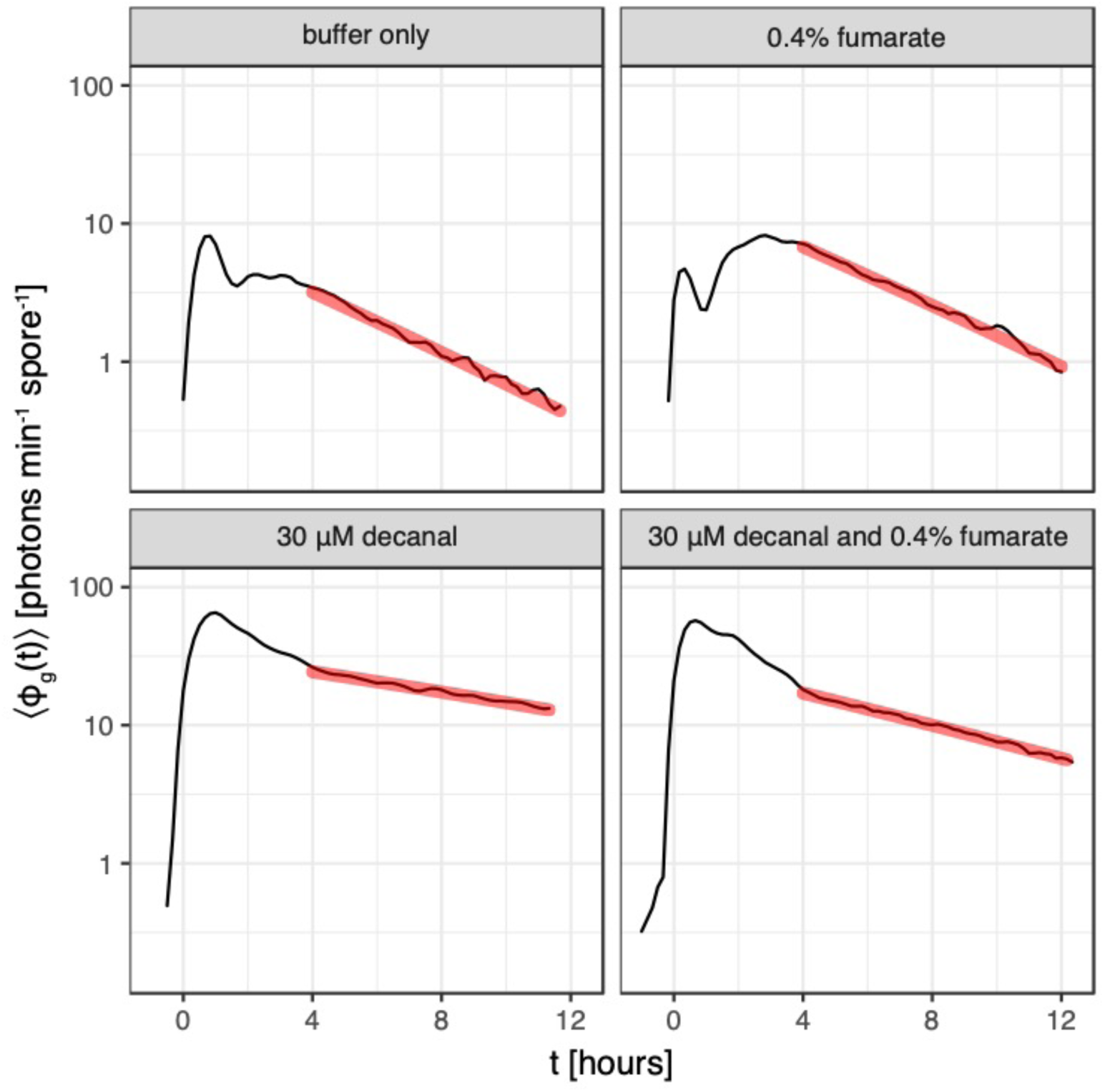
Bioluminescence signal decays exponentially after 4 h post-germination. The population-averaged bioluminescence dynamics of germinating spores ⟨*ϕ*_g_(*t*)⟩, depicted by the black line, exhibit exponential decline, beginning at 4 h post-germination. The rate of exponential decline *r* was estimated in each of 4 conditions: *r*_buffer-only_ = 22% h^-1^, *r*_fumarate_ = 22% h^-1^, *r*_decanal_ = 8% h^-1^, *r*_decanal+fumarate_ = 12% h^-1^. Estimated fits are depicted by red lines.

**Figure S3.**
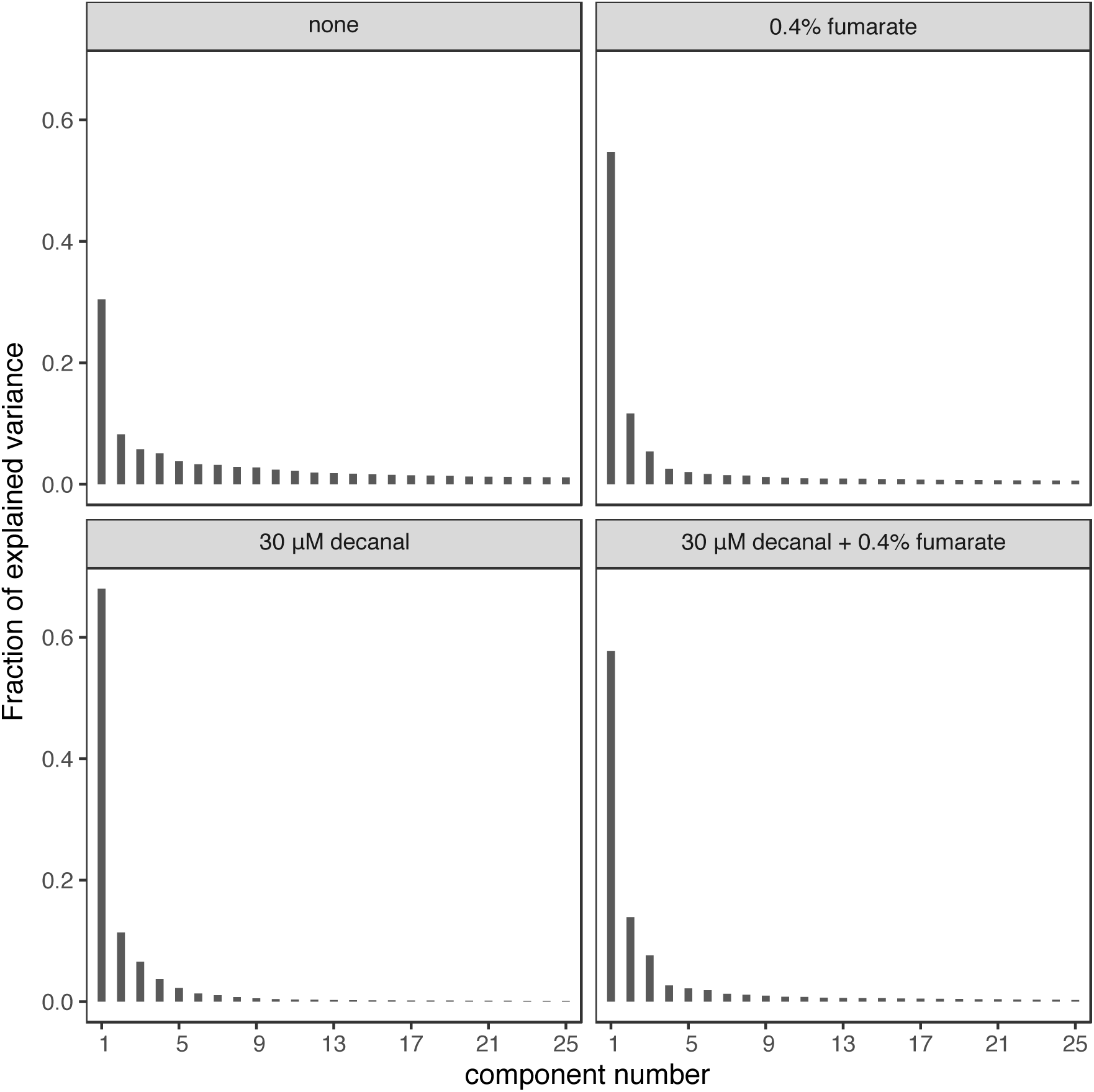
Variance explained by principal components. Principal component analysis was performed on the bioluminescent photon counts *ϕ*_g_(*t*) for each of 4 experimental conditions, as described in the Results section. The variance explained by principal component number *k* is *λ*_*k*_, and the fraction of variance explained is *λ*_*k*_(Σ_*i*_ *λ*_*i*_)^-1^. The data sets included 87 time points and therefore 87 principal components; only the first 25 components are included in the plots.

**Figure S4.**
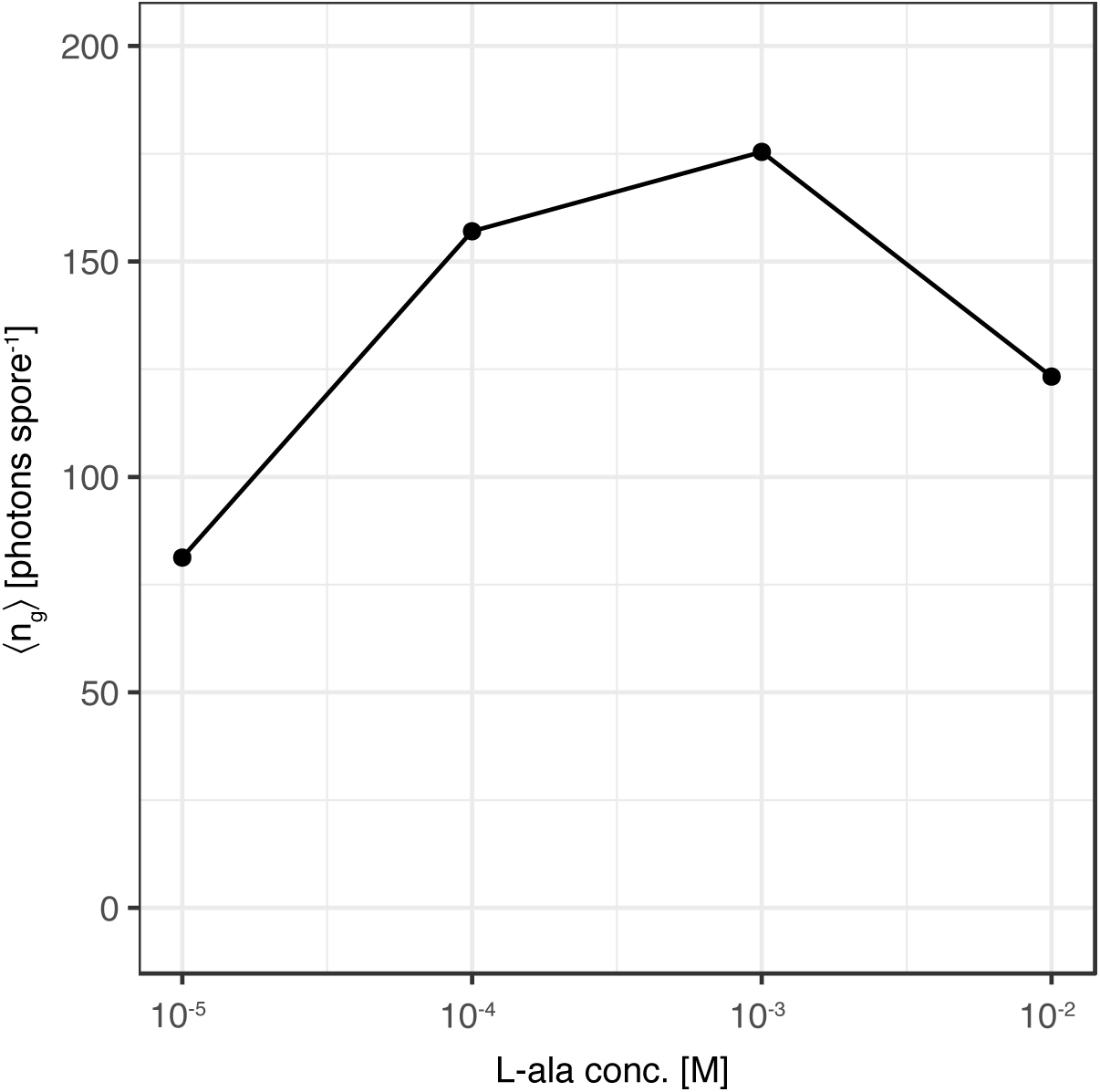
Bioluminescent photons counted per germinating spore. The bioluminescence signal as a function of time since germination, *ϕ*_g_(*t*) = *ϕ*(*t*-*τ*_g_), was integrated from 0 to 12 h post-germination to calculate the number of photons counted per germinating spore, *n*_g_. The average of *n*_g_ over all germinating spores is plotted against L-alanine concentration.

**Figure S5.**
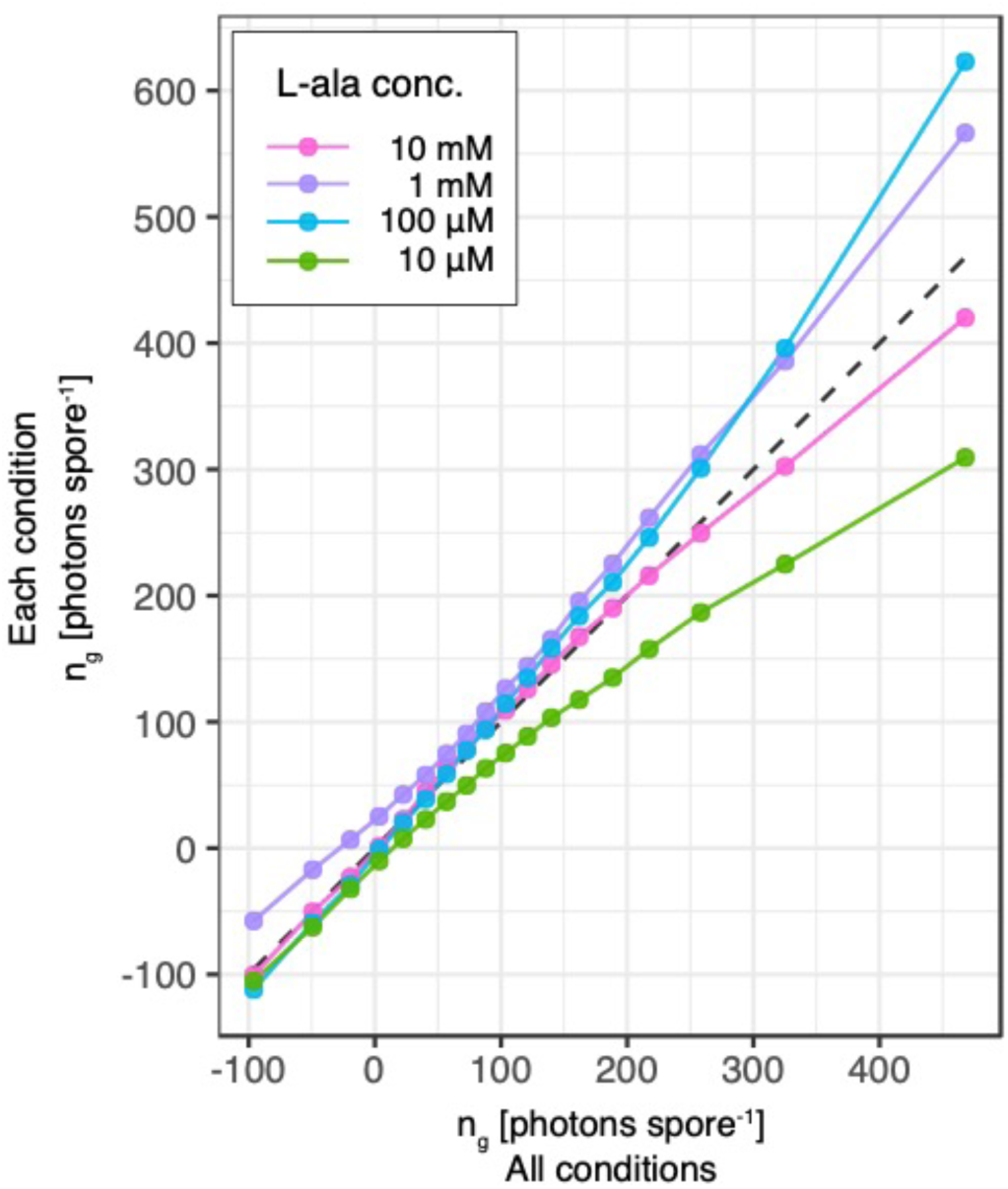
Quantile-quantile plot of bioluminescent photons counted per spore, germinating in response to 4 concentrations of L-alanine. Quantiles of the number of photons emitted per germinating spore during the 12 h post-germination, *n*_g_, were determined for 4 concentrations of L-alanine. Quantiles were also calculated for the data from all 4 L-alanine concentrations, weighted for equal contribution from each condition. The quantiles in each condition are plotted on the vertical axis, as a function of the corresponding quantile in all conditions. Quantiles were determined in increments of 5%, starting at 5% and ending at 95%.

